# Serial-passage assessment shows no confirmed resistance development to Norway spruce (*Picea abies*) resin in bacterial species relevant to wound infection

**DOI:** 10.64898/2026.05.08.723837

**Authors:** Kamilla Yamileva, Simone Parrotta, Maryam Ghanbarirad, Evgen Multia

## Abstract

The search for antimicrobials with a low propensity to select resistance has intensified in response to the global antimicrobial resistance crisis. Norway spruce resin (*Picea abies*) has long been used in Northern European wound care traditions and has shown broad antimicrobial activity in earlier microbiological studies. In the present study, we evaluated whether prolonged exposure to medical-grade spruce resin promotes reduced susceptibility in clinically relevant bacterial species. A 20-day serial-passage experiment was performed with *Staphylococcus aureus, Pseudomonas aeruginosa*, and *Enterococcus faecalis* using sub-inhibitory resin concentrations and broth microdilution readouts at baseline, day 10, and day 20. Resistance development was predefined as a ≥4-fold increase in inhibitory concentration. Baseline inhibitory concentrations were 1.25% for *S. aureus*, 5.0% for *P. aeruginosa*, and 2.5% for *E. faecalis*. After 20 days, inhibitory concentrations were 2.5%, 10.0%, and 2.5%, respectively, corresponding to at most 2-fold changes and remaining below the predefined threshold for resistance development. Validation and vehicle-control arms indicated that these shifts were not attributable to medium transfer or solvent-related bias. These findings suggest that medical-grade Norway spruce resin has a low short-term tendency to select for reduced susceptibility under serial-passage conditions.

**Graphical Abstract:** 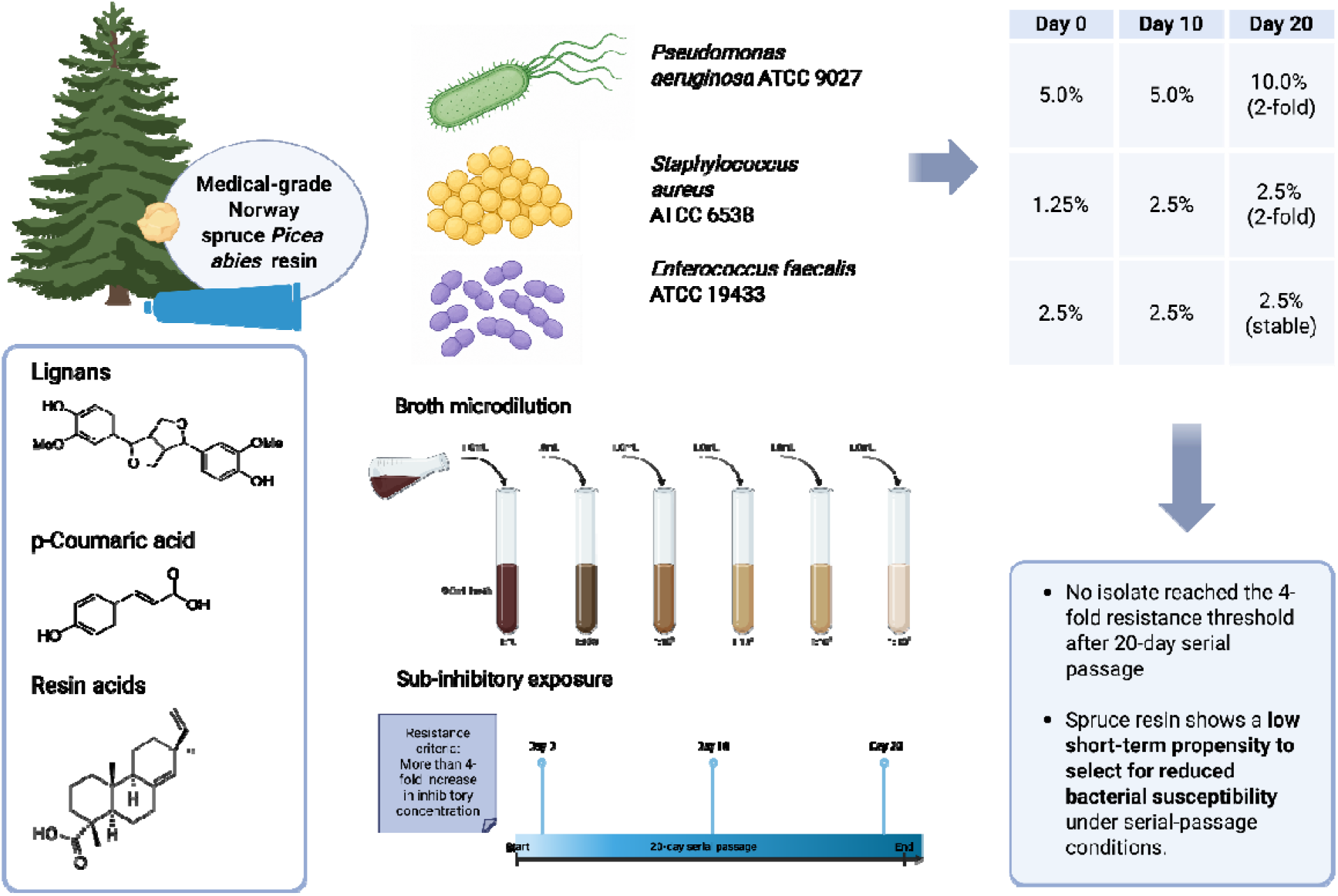

## 1. Introduction

Antimicrobial resistance has intensified the search for nonconventional antimicrobial agents that can control clinically relevant bacteria while exerting lower selective pressure for resistance than classical antibiotics [1, 2]. This need is particularly acute in wound care, where persistent microbial colonization, biofilm formation, and multidrug-resistant pathogens can complicate treatment and delay healing [3]. In some countries, topical antibiotic creams are still used in wound care [4], but routine use is discouraged because clinical benefit is limited, prolonged exposure may favor resistant bacteria, and repeated application can cause contact allergy [5]. In this context, plant-derived antimicrobials have attracted renewed interest because their activity often reflects chemically diverse phytoconstituents acting through multiple antibacterial mechanisms rather than a single defined target [6].

Among such natural antimicrobials, Norway spruce resin (*Picea abies*) has a distinctive position. In Northern Europe, salves prepared from spruce resin have been used in folk medicine for centuries to treat skin wounds, ulcers, and infections [7]. This long-standing traditional use of spruce resin-based wound products has inspired also modern laboratory [8– 10] and clinical investigation [11, 12]. Contemporary studies show that spruce resin and medical-grade spruce resin preparations have antimicrobial activity [7, 13–16], wound-healing potential [11, 12], and favorable topical safety profile [7, 17]. In the earliest modern *in vitro* work, home-made spruce resin salve showed bacteriostatic activity against clinically relevant wound-associated bacteria, with the clearest effects against Gram-positive organisms and demonstrable inhibition of methicillin-resistant *Staphylococcus aureus* (MRSA) and vancomycin-resistant enterococci (VRE) [13]. Later European Pharmacopoeia challenge-test studies extended this picture by showing broad microbicidal activity of the resin against *S. aureus*, MRSA, *Escherichia coli, Pseudomonas aeruginosa, Klebsiella pneumoniae, Bacillus subtilis*, and *Candida albicans*, with 10% resin in the salve medium sufficient to prevent microbial survival under challenge-test conditions [14]. Importantly, these studies also showed that apparent antimicrobial activity depends on assay format, since resin is poorly water-soluble and diffusion-based assays may therefore underestimate its effects, particularly against Gram-negative bacteria and yeasts [13, 14]. In chemical terms, spruce resin is not a single active substance but a complex natural mixture in which resin acids predominate, together with lignans and *p*-coumaric acid, a compositional profile that may help explain its broad biological activity [7, 14]. The antimicrobial activity of spruce resin likely arises from the combined effects of multiple constituent groups rather than from a single defined active compound, suggesting a mode of action that is broader and less target-specific than that of conventional antibiotics.

Mechanistic evidence further suggests that spruce resin does not behave like a classical single-target antibiotic. Ultrastructural and electrophysiological studies in *S. aureus* have shown cell-wall thickening, cell aggregation, altered fatty-acid composition, and dissipation of membrane potential after exposure to resin, resin salve, or abietic acid, indicating that antibacterial activity is associated with damage to the bacterial cell envelope and interference with energy metabolism [15]. It has been proposed that terpenic resin acids are the main antimicrobial mediators, acting through nonspecific membrane- and cell wall-associated effects, including disruption of the proton gradient and possible uncoupling of oxidative phosphorylation [7]. However, a recent study [18] of an aqueous spruce resin extract also showed marked ultrastructural damage in exposed microbes despite the absence of detectable non-oxidized resin acids and the presence of only low levels of oxidized resin acids, lignans, and *p*-coumaric acid, suggesting that antimicrobial activity may result from the combined effects of multiple resin-derived constituents.

In the spruce resin literature, the emphasis has so far been on antimicrobial spectrum, assay-dependent efficacy, ultrastructural mechanism, wound healing, biofilm eradication, and clinical use [7, 13–15, 19, 20]. In a recent case report, continuous bacteriological sampling of the wound of one patient was performed, and despite targeted intravenous antibiotic treatment, *P. aeruginosa*, ESBL-producing *K. pneumoniae, Morganella morganii*, and *E. faecalis* were cultured in the wound. One week after the use of spruce resin salve, it was observed that the pathogens cultured had changed to *Staphylococcus epidermidis, E. faecalis*, and *M. morganii [20]*. More recent studies have further extended the evidence base by showing that spruce resin-based salve can effectively eradicate mixed *P. aeruginosa*–*S. aureus* biofilm and penetrate its layers, even when the clinical isolates used exhibit high tolerance to bacitracin, neomycin, and polymyxin B [19]. However, these studies have not directly examined whether repeated sub-inhibitory exposure can select for reduced susceptibility. Thus, although spruce resin-based preparations have been considered less likely than conventional antibiotics to promote resistance, this question has not yet been tested in a controlled serial-passage experiment using medical-grade spruce resin. This question is relevant to chronic wound care, where microbial populations can change over time under the influence of environmental contamination, host-related selective pressures, and repeated antimicrobial treatment. Antibiotic exposure may favor less susceptible organisms, while biofilm growth can further increase tolerance and impair healing.

In the present study, we applied a serial-passage approach to medical-grade spruce resin used in resin salves such as Abilar, ilon Wundxtra Salbe, and SutriHeal Forte, to determine whether prolonged sub-inhibitory exposure can drive reduced susceptibility. *S. aureus, P. aeruginosa*, and *E. faecalis* were selected as clinically important bacterial species relevant to wound infection and commonly associated with antimicrobial resistance and reduced treatment responsiveness [13, 19, 21, 22]. Accordingly, this study aimed to determine whether repeated exposure to medical-grade Norway spruce resin makes bacteria less susceptible to its antimicrobial effects.

## 2. Materials and methods

### 2.1 Study design

A serial-passage resistance study was conducted to evaluate whether prolonged exposure to spruce (*Picea abies*) resin could lead to reduced bacterial susceptibility. The resin was manually collected from sustainably managed forests in Finland, purified according to patented method [23], and solubilized in ethanol. Serial dilutions were prepared in Mueller– Hinton broth (Biolife Cat No: 4017412) to final test concentrations of 10.0%, 5.0%, 2.5%, 1.25%, 0.625%, 0.3125%, and 0.15625%.

### 2.2 Test microorganisms and culture conditions

The organisms tested were *Staphylococcus aureus* ATCC 6538, *Pseudomonas aeruginosa* ATCC 9027, and *Enterococcus faecalis* ATCC 19433. Mueller–Hinton broth (MHB) was used for growth and Tryptic Soy Agar (TSA) (Neogen, Ref No: NCM0004B) for plating. Incubation was carried out at 30–35°C for 18 h.

### 2.3 Serial-passage protocol

This study was designed in accordance with CLSI guidelines for antimicrobial susceptibility testing (M07-A9: Methods for Dilution Antimicrobial Susceptibility Tests for Bacteria That Grow Aerobically) and adapted for resistance evaluation through serial passage. In this experimental setup, the test product was introduced at sub-inhibitory concentrations (typically 0.25× to 0.5× of the initial minimum inhibitory concentration (MIC)) into liquid culture medium. The selected test microorganism was inoculated and incubated under optimal growth conditions. After each incubation cycle, cultures were transferred daily into fresh medium containing the same sub-inhibitory concentration of the product, for a predefined number of passages (10 and 20 passages). MIC determinations were performed at baseline and at scheduled intervals (midpoint of 10 days and final passage after 20 days) using broth microdilution methodology. A ≥4-fold increase in MIC compared to the initial value was considered indicative of potential resistance development. This approach allowed monitoring of adaptive changes in susceptibility under prolonged exposure conditions. All assays were performed in duplicate. Since ethanol was used as solvent, vehicle controls containing ethanol without active resin were included. Validation passages without product exposure and validation MIC determinations from nutrient broth subcultures were also performed to exclude adaptation to laboratory conditions or medium transfer artefacts.

## 3. Results

### 3.1 Baseline inhibitory concentrations

At baseline (Day 0, Table 1), medical-grade spruce resin inhibited *P. aeruginosa* at 5.0%, *S. aureus* at 1.25%, and *E. faecalis* at 2.5%. Results were identical in the first and second trials, so a single combined summary table is shown. These values indicate that *S. aureus* was the most susceptible of the three tested organisms, while *P. aeruginosa* required the highest concentration.

**Table 1.** Summary of broth microdilution growth patterns for spruce resin during serial passage at Day 0, Day 10, and Day 20. All assays were performed in duplicate.

| Test organism | Time point | 10% | 5% | 2.5% | 1.25% | 0.625% | 0.3125% | 0.15625% | PC | NC |
| --- | --- | --- | --- | --- | --- | --- | --- | --- | --- | --- |
| *P. aeruginosa* | Day 0 | NG | NG | G | G | G | G | G | G | NG |
| *P. aeruginosa* | Day 10 | NG | NG | G | G | G | G | G | G | NG |
| *P. aeruginosa* | Day 20 | NG | G | G | G | G | G | G | G | NG |
| *S. aureus* | Day 0 | NG | NG | NG | NG | G | G | G | G | NG |
| *S. aureus* | Day 10 | NG | NG | NG | G | G | G | G | G | NG |
| *S. aureus* | Day 20 | NG | NG | NG | G | G | G | G | G | NG |
| *E. faecalis* | Day 0 | NG | NG | NG | G | G | G | G | G | NG |
| *E. faecalis* | Day 10 | NG | NG | NG | G | G | G | G | G | NG |
| *E. faecalis* | Day 20 | NG | NG | NG | G | G | G | G | G | NG |

**Abbreviations:** G = growth observed, NG = no growth observed, PC = positive control, NC = negative control. MIC endpoint was defined as the lowest resin concentration showing no visible growth. Resistance development was predefined as a $\geq 4$ -fold increase in MIC relative to baseline.

### 3.2 Serial-passage susceptibility over 20 days

Across 20 days of serial passage, susceptibility remained broadly stable. For *P. aeruginosa*, the inhibitory concentration remained 5.0% at day 10 and increased to 10.0% at day 20, corresponding to a 2-fold change. For *S. aureus*, the inhibitory concentration increased from 1.25% at baseline to 2.5% at day 10 and remained 2.5% at day 20, also a 2-fold change. For *E. faecalis*, the inhibitory concentration remained unchanged at 2.5% throughout the experiment.

Vehicle-control and validation arms (Table 2) remained stable over time and did not show progressive increases suggestive of methodological drift. The transfer procedure and medium changes did not appear to induce artificial resistance. Validation arms remained comparable to vehicle controls and did not show progressive increases over time. The day-20 increase seen in *P. aeruginosa* under test conditions was not mirrored in the validation arms. None of the tested bacteria showed a ≥4-fold increase in inhibitory concentration, and thus none met the predefined criterion for resistance development. The observed changes were limited to minor 2-fold shifts in *P. aeruginosa* and *S. aureus* (Tables 2 and 3).

**Table 2.**
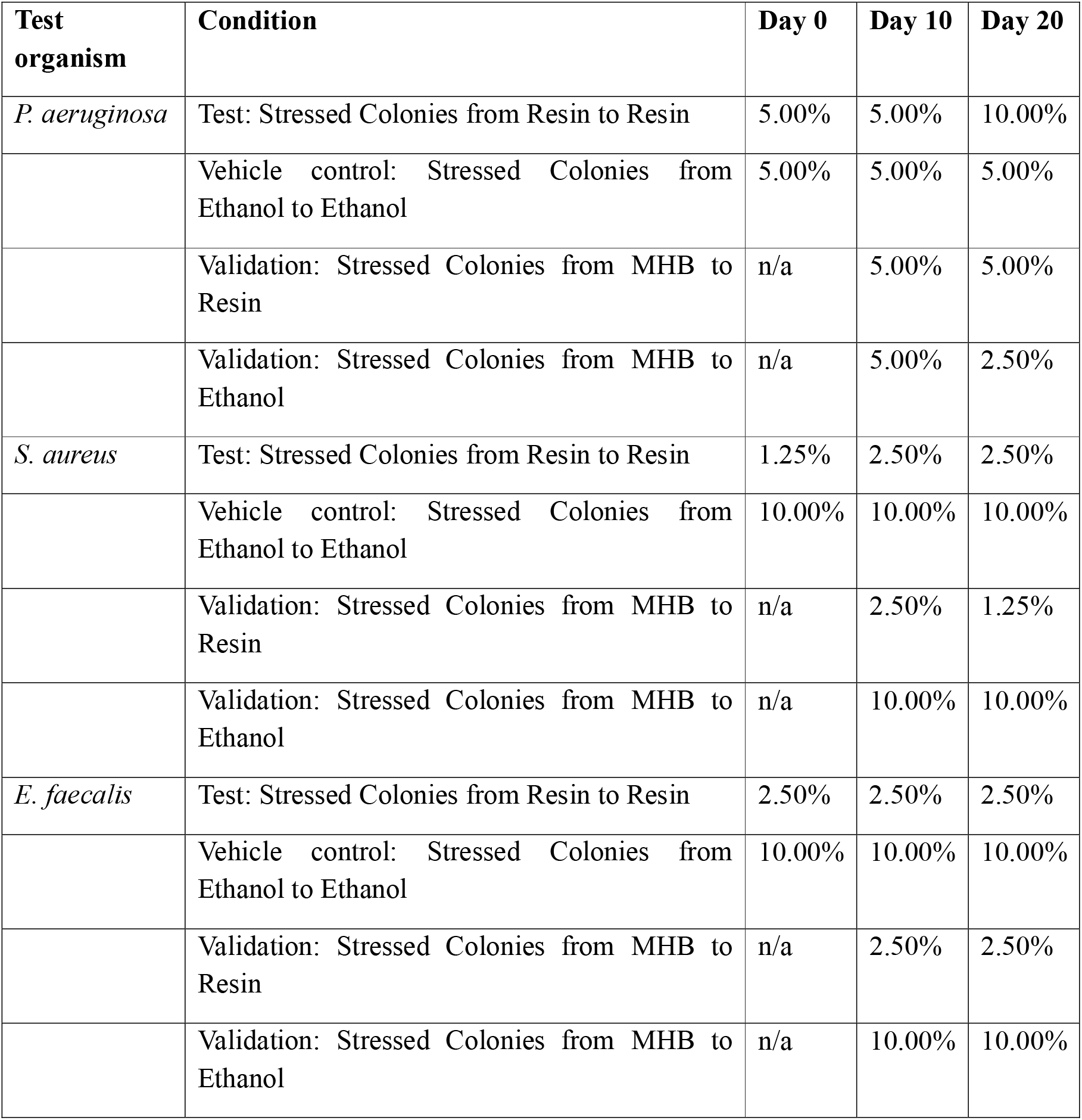
Validation and vehicle-control data across serial passage for spruce resin.

**Table 3.** Inhibitory concentrations of spruce resin during 20-day serial passage.

| Test organism | Baseline (Day 0) | Day 10 | Day 20 | Fold change Day 20 vs Day 0 | A ≥4-fold increase in inhibitory concentration resistance criterion met? |
| --- | --- | --- | --- | --- | --- |
| *P. aeruginosa* | 5.0% | 5.0% | 10.0% | 2-fold | No |
| *S. aureus* | 1.25% | 2.5% | 2.5% | 2-fold | No |
| *E. faecalis* | 2.5% | 2.5% | 2.5% | None | No |

20 days of serial passages were sufficient to assess short-term resistance selection potential because repeated sub-inhibitory exposure over many consecutive transfers gives bacterial populations sustained opportunity to adapt [24]. Overall, the susceptibility profiles remained stable, with only minor 2-fold variations in two organisms and no measurable change in the third.

## 4. Discussion

In this study, spruce resin showed a low tendency to select for reduced bacterial susceptibility during 20 days of serial passage in three clinically important wound-associated bacteria. None of the tested organisms reached the predefined resistance threshold of a ≥4-fold increase in minimum inhibitory concentration. Instead, the observed changes were limited to a 2-fold increase in *P. aeruginosa* and *S. aureus*, while *E. faecalis* remained unchanged throughout the experiment. These findings suggest that, under the present test conditions, repeated sub-inhibitory exposure to medical-grade spruce resin did not drive resistance-like adaptation. This has practical relevance for using spruce resin for infected wounds, where microorganisms may encounter repeated topical treatment under variable local exposure conditions, including sub-inhibitory levels at the wound surface or biofilm interface.

In serial-passage experiments, small changes, e.g., 2-fold shifts of *P. aeruginosa* and *S. aureus* do not necessarily indicate stable resistance [25, 26], especially when they remain below a predefined biological threshold and are not supported by consistent drift in validation or vehicle-control arms. The inclusion of ethanol controls and validation passages helped to exclude medium transfer effects, solvent-related bias, or simple adaptation to laboratory passage conditions. In particular, the day-20 increase seen in *P. aeruginosa* under resin exposure was not reproduced in the validation arms, supporting the view that the change was minor and not clearly indicative of progressive resistance selection.

Although closely comparable studies are scarce, published serial-passage experiments with conventional antibiotics provide useful context [27]. In MRSA, 50 serial passages to topical comparators such as mupirocin (64-fold increase), fusidic acid (256-fold increase), and retapamulin (16-fold increase) has produced MIC increases [27] well above the 4-fold threshold used in this study, and ciprofloxacin has similarly generated 32-fold MIC increases in both *S. aureus* and *P. aeruginosa* after 25 serial passages. Serial-passage selection has also been demonstrated in *E. faecalis* with linezolid [28]. These findings contrast with the 2-fold shifts observed for spruce resin in the present study, although differences in experimental design limit direct comparability.

These results extend earlier evidence of broad antimicrobial activity by addressing a question that previous spruce-resin studies had not directly examined, namely resistance selection under repeated exposure. Previous studies [13] demonstrated activity against Gram-positive wound-associated bacteria, including MRSA and VRE, and later challenge-test studies extended this evidence to Gram-negative organisms and *Candida albicans* [14]. Additional mechanistic studies suggested that spruce resin acts through relatively nonspecific effects on the bacterial cell envelope, including cell wall and membrane damage, altered fatty-acid composition, and dissipation of membrane potential [15]. Such a mode of action differs from that of classical single-target antibiotics [29] and may partly explain why marked resistance selection was not observed in the present serial-passage model. Differences between species were also biologically plausible. *S. aureus* was the most susceptible organism at baseline, whereas *P. aeruginosa* required the highest inhibitory concentration, consistent with the greater intrinsic tolerance often seen in Gram-negative bacteria and with earlier observations that apparent resin activity may depend on assay format [13, 14, 30]. *E. faecalis* remained stable during passage. The results suggest that any adaptive response to spruce resin, if present, was limited and species-dependent under the tested conditions.

From a clinical and translational perspective, these findings strengthen the rationale for continued use of spruce resin as a topical antimicrobial for wound management. Microorganisms in chronic wounds may encounter variable and sometimes sub-inhibitory exposure to topical antimicrobials because wound exudate can reduce bactericidal efficacy under wound-bed conditions [31], while biofilms increase tolerance and impair antimicrobial effectiveness [32, 33]. Against this background, it is important to identify antimicrobial agents that remain effective without easily causing bacteria to become less sensitive to them. This applies both to traditional spruce resin salves and to medical-grade resin formulations used in wound care. The present findings extend earlier studies by indicating a low short-term tendency of spruce resin to select for reduced susceptibility under the conditions tested.

The 20-day serial-passage design does not exclude slower adaptive responses that might emerge during longer exposure. In addition, the study assessed phenotypic changes in inhibitory concentration only and did not address whether repeated exposure induced stable genetic adaptation, altered fitness, cross-tolerance to antibiotics, or changes in biofilm behavior. Future studies should therefore include longer serial-passage experiments, broader strain panels, and clinically derived resistant isolates, particularly MRSA, VRE-associated enterococci, and multidrug-resistant *P. aeruginosa*.

## 5. Conclusion

Norway spruce (*Picea abies*) resin maintained antibacterial activity throughout a 20-day serial-passage experiment against *P. aeruginosa, S. aureus*, and *E. faecalis*. None of the tested organisms showed the predefined ≥4-fold increase in inhibitory concentration required to indicate resistance development. These findings suggest that, under the conditions tested, medical-grade spruce resin has a low short-term tendency to select for reduced susceptibility. Broader and longer-term studies are needed to determine whether this pattern is maintained across diverse and clinically resistant strains.

## Data availability statement

All data generated or analyzed during this study are included in this published article.

## Acknowledgements

Stathopoulos Aggelis and Giorgios Lagiopoulos from QACS Ltd are thanked for their help with the study.

## CRediT authorship contribution statement

**Kamilla Yamileva:** Writing – original draft, Writing – review & editing, Visualization. **Simone Parrotta:** Writing – review & editing, Methodology, Investigation. **Maryam Ghanbarirad:** Writing – review & editing, Resources, Funding acquisition, Project administration. **Evgen Multia:** Writing – original draft, Writing – review & editing, Visualization, Supervision, Resources, Project administration, Methodology, Investigation, Funding acquisition, Formal analysis, Data curation, Conceptualization.

## Ethics statement

None required

